# Deep learning-based location decoding reveals that across-day representational drift is better predicted by rewarded experience than time

**DOI:** 10.1101/2024.02.01.578423

**Authors:** Kipp Freud, Nathan Lepora, Matt W. Jones, Cian O’Donnell

## Abstract

Neural representations of locational and spatial relations in the hippocampus and related brain areas change over timescales of days-weeks, even in familiar contexts and when behavior appears stable. It remains unclear how this ‘representational drift’ is driven by combinations of the passage of time, general experience or specific features of experience. We present a novel deep-learning approach for measuring network-level representational drift, quantifying drift as the rate of change in decoder error of deep neural networks as a function of train-test lag. Using this method, we analyse a longitudinal dataset of 0.5–475 Hz broadband local field potential (LFP) data recorded from dorsal hippocampal CA1, medial prefrontal cortex and parietal cortex of six rats over ∼30 days, during learning of a spatial navigation task in an initially unfamiliar environment. All three brain regions contained clear spatial representations which evolve and drift over training sessions. We find that the rate of drift slows for later training sessions. Finally, we find that drift is statistically better explained by task-relevant experiences within the maze, rather than the passage of time or number of sessions the animal spent on the maze. While previous research has focused on drift as a measure of the changes in spiking activities of units, here we examine drift as a measure of change in oscillatory activity of local field potentials; our approach of using of deep neural networks to quantify drift in broadband neural time series unlocks new possibilities for defining the drivers and functional consequences of representational drift.

## Introduction

Neural activity patterns associated with sensation, cognition, and action are often not rigid but instead undergo progressive changes over timescales of minutes to weeks — a phenomenon called representational drift^1^. Such plasticity in the activity patterns of populations of neurons can occur even without apparent changes in environment or behaviour^2^. These findings challenge classical models which consider the stability of the engram as the basis of memory’s persistence^3^.

Representational drift is observed across different primary sensory modalities in the brain. For instance, neural representations in visual cortex exhibit drift in human, rat, and mouse^3–6^, with evidence suggesting that deeper areas of the visual cortical hierarchy encode visual information more persistently^7^. Similarly, olfactory representations change over time, even when external stimuli remain constant^8,9^. Representational drift is also evident in multimodal spatial representations, which have been best characterized in hippocampus and interconnected cortical regions. Soon after initial exposure to a new environment, place fields develop in the activity of neurons in the hippocampus and related brain regions^10^, but undergo drift both within- and across days^11–14^.

As one illustrative example, Driscoll et al.^15^ designed a virtual reality task where mice navigated a T-maze; while the animals’ performance remained consistent, chronic two-photon calcium imaging showed that the neuronal representation in the posterior parietal cortex shifted significantly over weeks. Despite these neural changes, the mice’s behavior and task performance did not show measurable alterations.

The mechanisms of representational drift remain unclear, but may relate to turnover of proteins, synapses, and neurons. Computational studies have suggested that ongoing synaptic plasticity can act as a source of noise, resulting in changes in neuronal dynamics reminiscent of drift at the population level, and leading to downstream changes in representations of already stored patterns^16^.

The computational and adaptive purposes of representational drift, if any, also remain unclear. Is it merely an epiphenomenon, or does it serve a functional role in neural processing? Several hypotheses have been explored^17^, one of which stems from the common experimental finding that neurons can go from highly active to entirely inactive on subsequent days. This may be analogous to ‘dropout’, an effective training strategy used in artificial deep neural networks to increase generalizability of learned representations^18^. Other research has suggested that this drift may be a way to ‘time-stamp’ different events, which follows intuitively from the observation that different sets of hippocampal place cells, potentially cells born at different times (due to hippocampal neurogenesis), are active in an environment on different days^19,20^. Alternatively, drift may allow the brain to sample from a large space of possible solutions, decreasing the likelihood of forming sub-optimal representations which exist as local minima in the loss landscape^21^.

The dynamics of drift have been quantified at the level of single units, and vary with brain region. Linear classifiers trained to predict odorants on data from single units in the primary olfactory cortex degraded to chance levels after 32 days (though this could be slowed by daily exposure to the same odorant)^8^. In mouse hippocampal CA1, long-timescale drift occurs orthogonally to the representation of context in network space, allowing for consistent readout of contextual information across weeks^22^. Exercise appears to increase the rate of CA1 drift^23^. One key question is whether drift is a function of experience or time? Khatib et al. (2023)^12^ found that within-day drift of spatial representations in dorsal CA1 is driven primarily by active experience rather than time. However Geva et al. (2023)^24^ examined drift across days, and found a more nuanced relationship — active experience in a maze drove changes in spatial tuning of dorsal CA1 cells, while time drove changes in the activity firing rates of cells. Importantly, both these studies equated ‘experience’ with ‘time spent on the maze’. This leaves open the question of whether different aspects of the animal’s task experience (time doing task, number of task trials, rewards) may have different impacts on representational drift^25^.

One other key mechanism modulating hippocampal representations over extended experience is ‘replay’, the reactivation of place cell activity patterns during sharp-wave ripples (SWRs) during resting and sleep states. Such replay may be implicated in representational drift as a signature of memories being integrated into new sets of neurons as memories are replayed and consolidated^26^. Consistent with this, Grosmark et al. (2021) found that replay during SWRs was correlated with the long-term stability of spatial representations within single units in the hippocampus^27^.

Here, we develop and test methods to disentangle the impacts of time, experience, and SWR-associated replay on representational drift tracked simultaneously across multiple brain regions and ethologically-relevant timescales of 2–3 weeks. We recorded local field potentials (LFPs) simultaneously from chronically-implanted adjustable microelectrodes spanning three distinct brain regions of six rats; dorsal CA1 of hippocampus (CA1), prefrontal cortex (PFC), and the parietal cortex (PC).

Our approach^28^ uses deep convolutional neural networks (CNNs) trained on wavelet decomposition images of this LFP data^29^ to decode animal location in a multi-session longitudinal dataset. This offers several key advantages compared to traditional spike-based analyses. First, most spike-based methods rely on the assumption that the firing activity of neurons is independent, which is a problematic simplification when trying to quantify representational drift at the ensemble level. Neural activity is highly interconnected, and spiking neurons often exhibit correlated activity, a feature that is difficult to capture without modeling pairwise or higher-order correlations. While it is possible to construct decoding models that include these correlations, fitting such models to real data is computationally intensive and becomes increasingly complex as the number of neurons grows^30^.

Spike-based approaches also require spike sorting, a process where raw extracellular signals are analyzed to detect action potentials (spikes) and assign them to individual neurons. This is often done by analyzing the shape of the spike waveform, which can be affected by factors such as electrode placement and noise. Spike sorting is computationally demanding, technically challenging, and relies on algorithmic decisions that may discard valuable information in non-spike frequency bands^31^. As a result, traditional spike sorting focuses primarily on discrete spiking events, ignoring the wealth of information contained in lower-frequency oscillations. In contrast, our method, based on raw LFP data and wavelet decomposition, captures oscillatory dynamics across a broader frequency spectrum, offering a more expansive view of neural activity that includes slow and fast oscillations. Additionally, it is particularly difficult to monitor the same population of neurons over many days across multiple brain regions, adding another layer of complexity to performing spike-based analyses on longitudinal datasets.

Neural oscillations are rhythmic patterns of electrical activity generated by the interactions between excitatory and inhibitory neurons within the brain; they reflect the coordination of large populations of neurons and provide insight into global brain dynamics, including rhythmic processes related to memory, attention, and cognitive states^32^. These oscillations are present across various frequency bands, such as delta, theta, gamma, and beta, each associated with different cognitive and behavioral functions. For example, theta oscillations are often linked to navigation and memory processes, while gamma oscillations are involved in attention and sensory processing. Oscillations emerge as a result of the synchronized firing of neurons, which together form larger-scale patterns of brain activity, giving researchers a window into how different brain regions communicate and synchronize over time^33,34^.

The deep convolutional neural networks (CNNs)^35^ used in our approach are particularly suited to analyzing these oscillatory dynamics because they do not require spike sorting, allowing for minimally processed input. Using CNNs to decode behavioural variables from wavelet decompositions of intercranial EEG data was first performed by Frey et al. (2019)^36^, who achieved state of the art location decoding results using broadband data recorded at 15000 Hz from 128 hippocampal recording sites. We use an equivalent wavelet-based method on our lower frequency recordings, which transforms our LFP data into a time-frequency representation, effectively capturing oscillations across different frequency bands without focusing on spiking events. This is crucial because we aim to explore drift in terms of changes in oscillatory patterns rather than spikes, providing a broader view of neural dynamics. This method ensures we capture the time-varying nature of these oscillations, which is important for tracking representational drift over time.

By focusing on oscillations, our method offers a complementary perspective to spike-based analyses of representational drift. Spiking events are inherently sparse and may not fully capture the ongoing network-level processes that contribute to drift. In contrast, oscillations provide a continuous measure of neural activity and are sensitive to the dynamic interactions between different brain regions. Importantly, the wavelet decomposition method we employ captures time-varying frequency components, enabling us to track how oscillatory activity evolves over time, independent of individual spikes. This approach could also be applied to datasets such as scalp EEG or MEG, which are not well-suited for spike-based analyses, offering broader translational potential for human drift-analysis experiments^37^.

## Results

### Hippocampal, parietal and prefrontal LFPs code spatial representations that become more informative with behavioural training

We recorded local field potential data from hippocampal CA1 (HPC), prefrontal cortex (PFC), and parietal cortex (PC) from 6 rats as they navigated a memory and decision-making task to find sucrose rewards on a 118 *×* 164 cm maze. Before data collection, the rats underwent three sessions of passive exploration in the maze to familiarize themselves with its layout. Recordings were then conducted one session per day for approximately 30 days (Figure 5)^38,39^. During the initial phase of training, the rats were trained using an “alternate turn” rule, requiring them to navigate in a direction opposite to their initial guided turn (i.e., if the rats were directed to turn right at the guided turn, they should then turn left at the choice point). Once task competency was achieved (defined as three consecutive sessions with at least 85% of trials correct), the rule was reversed to a “match turn” rule, where rats were rewarded for turning in the same egocentric direction as their initial guided turn. Throughout these experiments, rat location was recorded via head-mounted LEDs.

We first trained deep CNNs to decode rat maze location using 0.5–475 Hz wide-band LFP data collected from the hippocampus (HPC), prefrontal cortex (PFC), and parietal cortex (PC). A separate model was trained for each combination of rat, experimental session, and brain region. After training, we evaluated the location decoding performance using a withheld testing set of unseen data. All decoders performed significantly better than random, with the HPC-only models achieving an average location decoding error of 48.32 cm (on a maze measuring 164 × 118 cm). Decoders based on PFC and PC data resulted in slightly higher errors, averaging 50.08 cm and 51.38 cm, respectively (Figure 1B).

**Figure 1.**
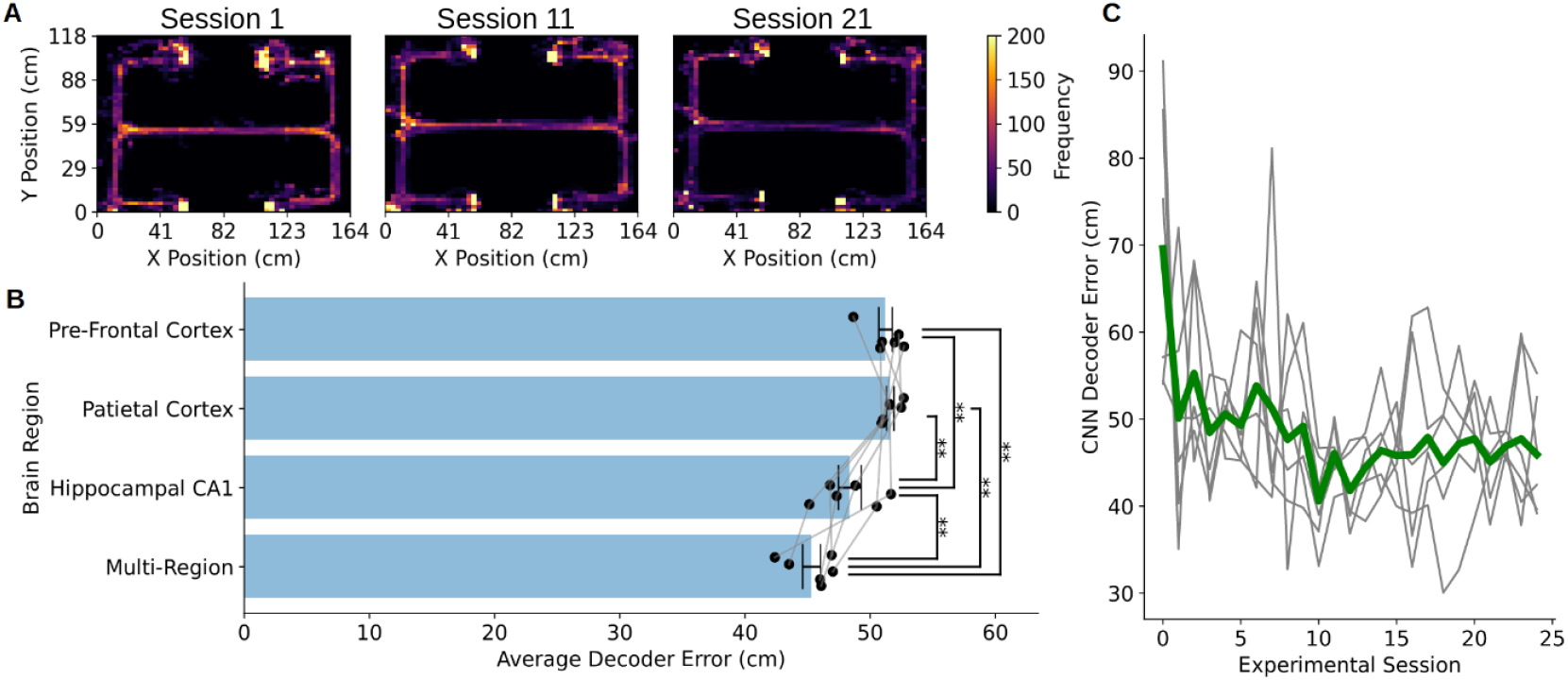
(A) Heat maps showing the location of a rat throughout the experiment - we note that although the experimental protocol and the performance of the rat was changing, it’s location distribution remains the same. (B) Neural networks which were trained on different brain regions obtain different average error, with models trained on hippocampal CA1 data performing significantly better than models trained on data from other brain regions, and a multi-region model performing significantly better than every single region models (“**” indicate significance at p<0.05, ANOVA with Tukeys correction). (C) Decoder error decreased as training session number at which the data was collected increased. Grey represents decoder error for individual rats, while green is mean decoder error of a neural decoder (each trained on data from one session) for all rats.

While these decoding errors are higher than those typically achieved using broadband LFP data that include high-frequency, spike related features^29^, it is important to note that our models were trained exclusively on the oscillatory activity present in the wavelet decomposition of the LFP signal. Crucially, we did not include spiking information, which is known to carry more precise location data^37^. This is a deliberate aspect of our method, as our primary focus is on understanding representational drift in oscillatory activity, not on spike-based drift. Therefore, the relatively higher decoding errors reflect the use of purely oscillatory dynamics, which are inherently less precise than spiking data for tasks such as location decoding. However, this focus on oscillations provides a complementary perspective to traditional spike-based analyses, allowing us to explore how changes in the rhythmic coordination of neural populations may drive drift over time.

While we observed that our decoding errors were higher than those reported in spike-based approaches, the errors did increase significantly with train-test-lag (the time difference between training and testing data) across all brain regions considered (Figure 2). This consistent increase in error suggests that our system successfully detected drifting spatial representations in the neural data over time. The progressive divergence in decoding accuracy indicates that the neural activity patterns responsible for encoding spatial information were not static but instead changed gradually, contributing to the observed representational drift.

**Figure 2.**
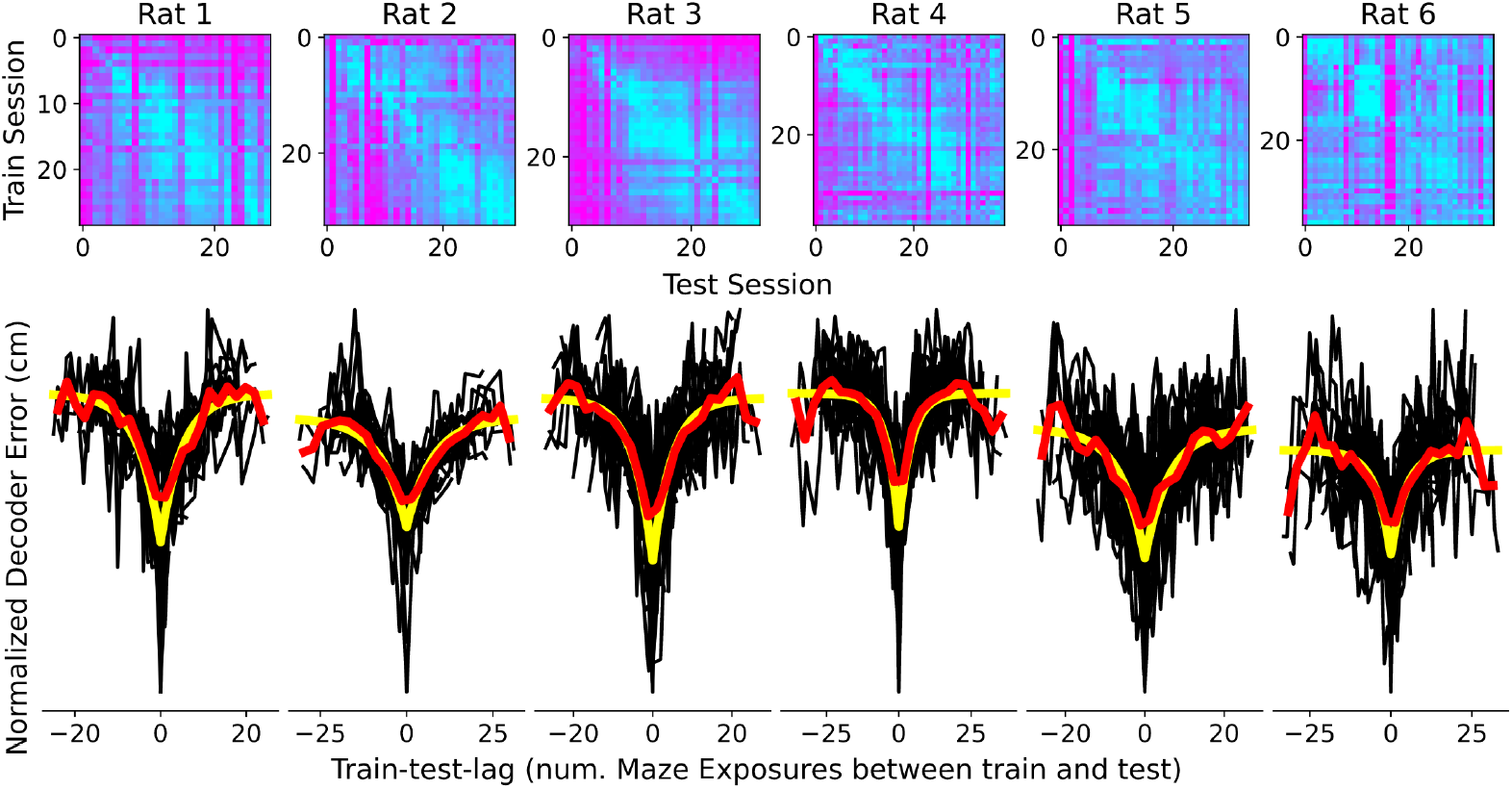
Above: Accuracy table for all rats, with each subsequent row representing a later session of training data, and each subsequent column representing a later session of testing data. The diagonal represents error with zero train-test-lag i.e. when training and testing on different data collected from the same day. Below: Decoder error against train-test lag for the multi-region models of all rats; the mean values of these error curves is shown in red, and fitted exponential decay curves are also shown in yellow.

This pattern of increasing error with time-lag reinforces the idea that our method is sensitive to temporal changes in oscillatory dynamics and the gradual evolution of spatial representations. Importantly, while the absolute decoding errors were higher compared to methods using spiking information, this level of accuracy is sufficient for answering our primary questions about representational drift in oscillatory activity. Although the rats were learning to improve their performance in the memory and decision making task during the sessions, they were not learning the spatial layout of the maze itself, as they had already been fully familiarized with the environment through passive exploration prior to data collection. As our focus is on decoding location within a maze that was already well-known to the animals, it remains viable to quantify drift in spatial representations in this way. We note that the goal of this analysis is not to achieve the most precise spatial decoding, but rather to quantify how spatial representations evolve over time and experience, as reflected in changes in the oscillatory behaviors in LFP signals. Since we aim to understand drift in oscillatory activity rather than spike-based location encoding, these results align well with our research objectives.

Next, we trained deep CNNs to decode location using LFP data from all three brain regions simultaneously (HPC, PFC, and PC), with one model trained for each rat and experimental session combination. By leveraging information from multiple brain regions, these models achieved an average decoding error of 48.63 cm, significantly outperforming both random decoding and the single-brain-region models (Figure 1B). The combined data provided a more comprehensive picture of neural activity, allowing for more accurate location decoding compared to using individual regions in isolation.

Additionally, we found that the mean decoder error decreased as the rats progressed through the initial training sessions. Specifically, during the first 10 experimental sessions, decoding accuracy improved across all rats, with a clear and significant reduction in error (Figure 1C). A regression coefficient of -0.9782 and a p-value of 0.021 was calculated, indicating a significant relationship between session number and error reduction during this early phase of training. However, after the 10th session, no further significant decreases in decoding error were detected, suggesting that the neural representations had stabilized by that point.

To test whether low- vs high-frequency bands carried more location information, we trained separate models using LFP data either above or below 300 Hz. This frequency was selected as it split our available frequency bands in half, and groups together all commonly discussed oscillatory activity in the lower frequency bands, from slow delta oscillations to sharp wave ripples, while grouping together all higher frequency bands potentially containing spiking information. The results showed that the vast majority of location information was contained in the lower frequency range (below 300 Hz), with models trained on higher frequency data performing only marginally better than random at same-day decoding. In contrast, models trained on the lower-frequency bands (*<* 300 Hz) performed almost as well as models trained on the full frequency range.

This is consistent with known neural oscillatory dynamics. The lower frequency bands, which include theta (4–12 Hz), gamma (30–100 Hz), and sharp-wave ripples (100–200 Hz), are implicated in cognitive processes such as spatial navigation, memory consolidation, and neural synchronization^32^. Theta oscillations, in particular, have been extensively linked to hippocampal function during navigation and memory tasks, while gamma oscillations are thought to facilitate communication between different brain regions. Sharp-wave ripples (SWRs), typically occurring in the 100–200 Hz range, play a key role in replaying and consolidating memories during sleep or quiet rest immediately following rewards. The strong location information carried by these lower-frequency oscillations aligns with their well-established involvement in coordinating cognitive functions across neural networks. Conversely, the higher-frequency bands (above 300 Hz) contain less location-specific information, which may explain the lower decoding performance when using only those bands.

### Oscillatory representations drift over time

Using results from both the multi-region models (trained on data from all brain regions) and the single-region models, we observed three key findings. First, for all rats and brain regions, decoding error increased consistently as the train-test lag increased (Figure 2). This observation indicates significant representational drift over time, even when analyzing only nonspiking lower-frequency LFP data. Statistical testing confirmed that this drift was highly significant across all regions (p *<* 1 *×* 10^−5^ for multi-region, CA1, PFC, and PC models).

Second, the relationship between mean normalized accuracy and train-test lag followed a pattern best described by exponential decay curves:

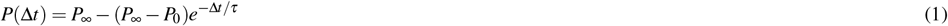

where

- Δ*t* represents the train-test lag.
- *P*_0_ represents decoder performance when the train-test lag is zero (i.e., when training and testing data are from the same session).
- *P*_∞_ represents decoder performance at very long train-test lags.
- *τ* is the decay time constant, which quantifies how quickly the decoding error approaches its asymptotic value as the train-test lag increases. Specifically, *τ* is the train-test lag at which decoder error will rise 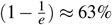 of the way to its asymptotic value

In this model, a small value of *τ* indicates faster drift, where the representations change rapidly over time, whereas larger *τ* values suggest slower drift, where the representations remain more stable across sessions. Based on these curve fits, we found that the mean decay time constants (*τ*) ranged between 5–12 sessions for individual rats. Interestingly, the *τ* values did not differ significantly between the brain regions considered (Figure 3C; p = 0.445, ANOVA), suggesting that the drift rates were comparable across hippocampal, prefrontal, and parietal regions.

**Figure 3.**
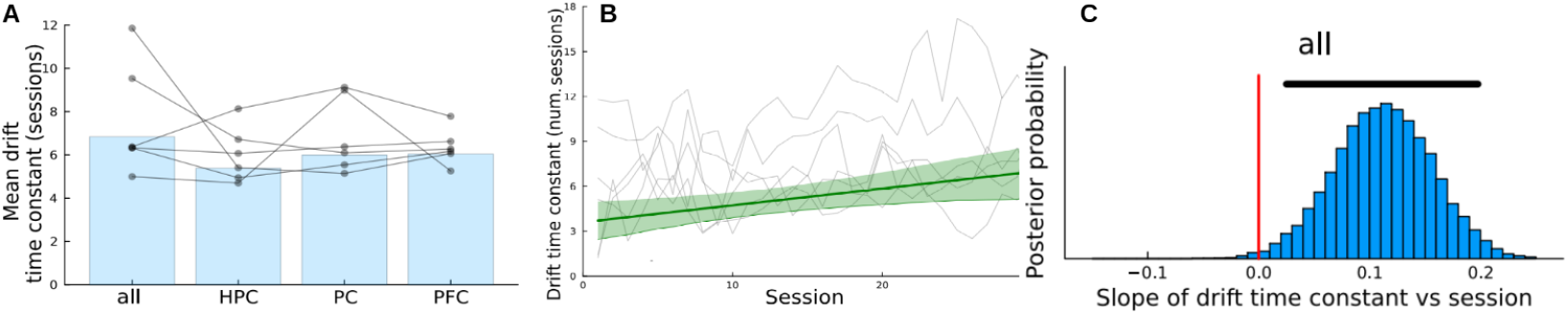
(A) Decay time constants for fitted exponential decay curves for models trained on different brain regions. (B) Decay time constants over time for the multi-region models of all rats, with the upwards trend indicating increasing stability. Posterior mean linear fit plotted in dark green with 95% highest density interval in light green. (C)Posterior probability of the slope parameter for a linear regression between drift time constant vs training session. Black bar indicates 95% highest-density interval.

### Oscillatory representations become more stable over time

To determine whether the rate of representational drift changed as training progressed, we applied a Bayesian analysis (see Methods) to calculate the drift time constant *τ* for each individual training day. Each row in the matrices shown in Figure 2 represents one training day, allowing us to track how the drift time constant evolved over time. For models trained using multi-region data, we observed a clear trend: the drift time constant increased on average as the number of training sessions increased (Figure 3). This was supported by the 95% highest density interval (HDI) for the slope parameter, which was [0.025, 0.198], indicating a very low probability (0.0064) that the slope was less than zero. This result suggests that as training progressed, representations became more stable, with slower drift.

For the single-region models, the evidence for increased drift time constants was weaker but still suggestive of the same trend. The 95% HDIs for the slope parameters were [-0.034, 0.15] for HPC, [-0.015, 0.165] for PC, and [-0.022, 0.138] for PFC, with the probability of the slope being less than zero at 0.0725, 0.05295, and 0.0984, respectively. Although these results are less conclusive than those for the multi-region models, they still hint at an overall increase in the stability of spatial representations in each of the three regions considered.

### Drift is a function of both replayed and active experiences rather than time

In the previous sections, drift curves, such as those shown in Figure 2, were ordered based on the number of experimental sessions, where points at x = *±*1 indicated that the train-test pair was separated by *±*1 experimental day. However, experiments were not always conducted daily; some single days or entire weekends were skipped. Moreover, not all experimental days contained the same number of trials, and rat performance varied from day to day. Additionally, varying numbers of hippocampal SWR events were detected during rest periods between experimental sessions. To better understand the causes of drift, we incorporated this variability to statistically examine how drift is influenced by different experimental factors.

To do this, we re-represented the train-test lag in terms of various experimental parameters:

1. The number of experimental sessions between the given train-test pair
2. The number of days between the given train-test pair
3. The number of guided trials which occured between the given train-test pair
4. The number of incorrect choice trials which occured between the given train-test pair
5. The number of correct (rewarded) choice trials which occured between the given train-test pair
6. The total number of choice trials which occured between the given train-test pair
7. The number of SWRs detected during rest periods between experimental sessions.

We then re-fitted exponential decay curves to these newly ordered data, assessing the fit between the exponential model and observed decoder performance error. To quantify the effect of each parameter, we normalized the error by dividing it by the mean error for each rat, allowing for comparison across variables. As shown in Figure 4, we observed a significant reduction in fitting error when the data was organized by experiential variables (e.g., trial numbers, performance) compared to ordering based solely on the time between sessions or the number of exposures to the maze.

**Figure 4.**
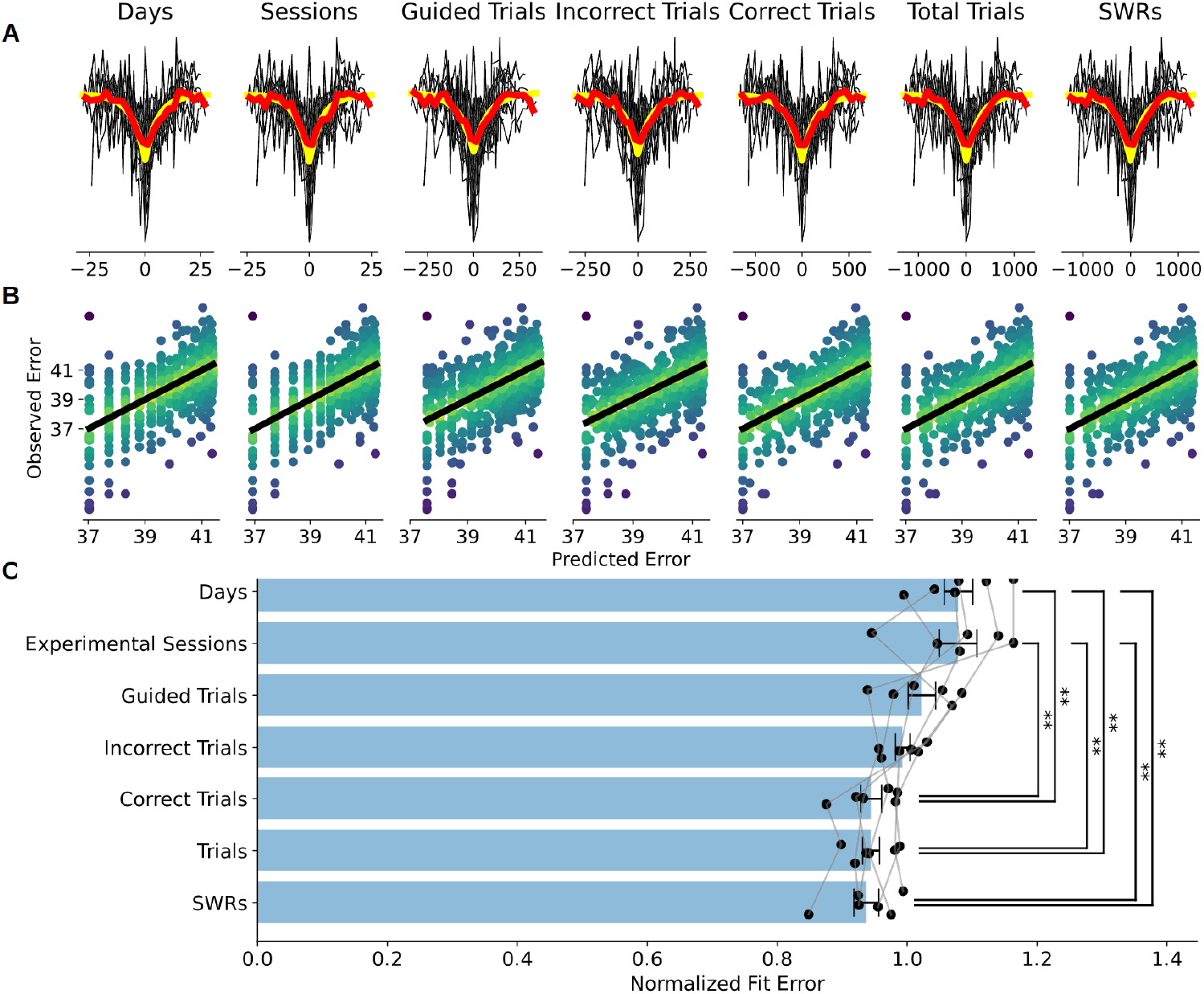
(A) Decoder error against train-test lag for one rat, with lag being represented by different experience- or time-related experimental variables. The mean values of these error curves is shown in red, and fitted exponential decay curves are shown in yellow. (B) Expected vs. observed prediction error for each of the train-test-lag fits presented in A. (C) Normalised error between observed error values and fitted exponential decay curves, when using each type of lag variable, normalized by the average of all lag variables. We note that when using certain experienced based variables as our train-test lag, we obtain significantly less fit error than when using time based lag, or using a simple exposure count based lag. We also that using the number of sharp wave ripples SWRs as our train test lag parameter results in the lowest fit error. “**” indicate significance at p<0.05, ANOVA with Tukeys correction.

Among experience-related variables, the largest reduction in error occurred when ordering by the number of total trials and, importantly, by the number of correct trials (i.e., trials where the animal was rewarded). This suggests that drift is more tightly related to rewarded experiences than simple exposure or time spent on the maze. Notably, we observed the largest reduction in error when organizing the data by the number of SWR events detected during rest between sessions. This highlights the significant role of replay during SWRs in driving representational drift, suggesting that both active task engagement and neural replay during rest are important for understanding the dynamics of drift.

## 1 Discussion

We have presented a method for the quantification of representational drift by using a deep convolutional neural network (CNN) trained on wavelet decomposition images of local field potential (LFP) data (Figure 7). This approach allowed us to measure representational drift without the need for either spike sorting, a time-consuming and computationally intensive process, or stable. multi-day recording of unit activity from multiple brain regions. Our results show that across-day drift is well characterized by exponential decay curves (Figure 2). Furthermore, we find that drift is better predicted by experience-related variables than by simple time-based measures. Notably, drift was most effectively predicted by replayed experiences, particularly by the number of sharp-wave ripple (SWR) events during sleep, underscoring the importance of replay in driving drift. Importantly, these findings relate specifically to oscillatory activity and not the more commonly studied spike-based representations.

Previous research has primarily focused on spiking data to study representational drift, but our work demonstrates that oscillatory dynamics—captured through local field potentials (LFPs)—also show a progressive stabilization over time. This finding highlights a new perspective on how the brain encodes and maintains spatial representations. Given that the rats in our experiment were actively learning the task during recordings, out results align with the hypothesis that representational drift is initially fast when exposed to a novel context but slows as the brain becomes more familiar with the environment and task demands. This phenomenon is consistent with other research suggesting that ensemble-level drift is more pronounced upon first exposure to a new context, gradually stabilizing as familiarity increases^40^. By focusing on the oscillatory components of neural activity, we provide a complementary perspective on drift, suggesting that changes in network-level oscillatory patterns contribute to representational instability. This is significant because oscillations represent the coordinated activity of large neural populations, providing insights beyond the single-neuron spiking examined in traditional drift studies.

Our results also demonstrate that models combining data from multiple brain regions significantly outperform models using single-region data (Figure 1B). This highlights the distributed nature of spatial information and the importance of interactions between brain regions in generating stable representations of space^41,42^. Our CNN-based approach effectively captures these inter-region relationships, whereas more traditional spike-based decoders often assume independence between neural firing, limiting their ability to account for these correlations^43^. This advantage of our approach underscores the value of examining non-independent neural dynamics, such as those seen in oscillatory activity.

Historically, early research on representational drift focused on the effects of time as the primary driver of drift dynamics^15,17^. More recent studies, using calcium imaging, have shifted the focus toward experience as a key factor driving drift^12,24^. While our findings are consistent with these studies, our analysis goes further by differentiating between types of experience, including both active engagement and putative experiences. We found that rewarded experiences and replayed SWRs were the strongest predictors of drift. These results are in line with recent findings showing that high reward expectation can slow the rate of drift, while reduced reward expectation speeds it up^25^. Our study extends these findings by showing that this relationship holds over a longer timescale (30 days) than the 4-day period previously examined.

Furthermore, our work is the first to explicitly link representational drift to hippocampal sharp-wave ripples (SWRs), suggesting that the reactivation of neural patterns during sleep plays a critical role in the stabilization or destabilization of spatial representations. This provides novel evidence that both active experiences (like task-related engagement) and replayed experiences (during SWRs) drive drift. This insight underscores the importance of considering replay mechanisms in understanding how representations evolve over time. SWRs, which have been linked to memory consolidation in previous studies^26^, are likely a key mechanism by which the brain reactivates and potentially modifies stored representations, thereby influencing the trajectory of drift.

The use of LFPs to study drift is another novel aspect of this work. Previous studies examining drift in LFP data have mainly focused on obtaining stable, low-dimensional representations rather than quantifying the dynamic changes in neural activity across days^44^. In contrast, most research on representational drift has focused on spiking activity at the single-neuron level^12,15,24^. However, LFPs provide a more comprehensive view of the aggregate neural activity within a region, allowing us to examine the network-level dynamics that contribute to representational drift. By focusing on oscillatory activity, we capture aspects of neural processing that are missed when focusing exclusively on spikes. This shift in perspective opens up new possibilities for understanding drift at both the local and global network scales.

Our findings suggest that oscillatory drift is a distributed process across multiple brain regions and frequency bands, with interactions between brain areas playing a key role in maintaining or reshaping representations over time. The multi-region models demonstrated significantly better performance than single-region models, underscoring the importance of inter-regional communication and coordination in the brain’s handling of spatial information. This distributed nature of drift emphasizes the need for future research to explore the specific roles that different brain regions and oscillatory frequencies play in modulating drift dynamics.

This oscillatory drift framework provides a complementary perspective to traditional single-unit spiking-based analyses. Although spiking activity offers valuable insights into how individual neurons contribute to memory and spatial representations, our use of LFP data uncovers the underlying oscillatory dynamics that are critical for understanding large-scale population activity. In particular, LFP oscillations like theta, gamma, and sharp-wave ripples are believed to facilitate inter-regional communication and memory consolidation, processes that are likely involved in drift. By focusing on the aggregate dynamics rather than individual neuron spikes, we provide a broader understanding of how network-level interactions and oscillations contribute to drift.

One strength of this approach is that it can be easily generalized to other brain regions or modalities of neural data. Our method, which decodes drift by quantifying changes in the performance of a CNN trained on wavelet images of LFP data, offers flexibility in its application. It could be adapted to other types of time-series data, such as calcium imaging or even non-invasive scalp EEG recordings in humans, to track representational drift in different contexts and systems. For example, calcium imaging data, which tracks the activity of large neural populations, could similarly be used to assess how representational drift unfolds over time, offering a way to link insights across spiking activity, LFPs, and other neural signals^29,45^.

Moreover, our results show that drift dynamics are most strongly predicted by reward-related experiences and replayed experiences (via SWRs), which highlights the importance of reward-driven learning and sleep-based consolidation in stabilizing representations. While previous studies have linked representational drift to ongoing task engagement and learning, the role of replay in modulating drift is less understood. Our findings extend this line of research by showing that SWRs, which are known to correlate with reactivation and consolidation of spatial memories during sleep^26,27^, play a key role in the dynamics of drift. This suggests that drift is not merely a consequence of ongoing neural plasticity but may also reflect the brain’s active processes of reinforcing or reshaping representations during restful states. Our findings align with emerging research that suggests neural plasticity and learning dynamics are not uniformly driven by time but are heavily influenced by behavioral context and task demands^25^.

In conclusion, this study presents a novel approach to understanding the intricacies of representational drift, showing that it drift affects not just spike-based representations, but oscillatory dynamics as well. While time and experience have been recurrent themes in prior studies, our findings imply that specific kinds of experiences, especially reward-related and replayed experiences, are drivers of ensemble coding dynamics. Further, our approach using LFP data as a modality for analyzing drift offers a complementary and promising avenue for future research. As we refine our understanding of oscillatory drift, we open the door to new questions about the role of network-level dynamics in memory, learning, and representation in the brain.

## Methods

### Data collection

Data was collected from 6 adult, male Long-Evans rats (300-500g, Harlan UK) navigating a memory and decision-making task to find sucrose rewards on a 118 *×* 164cm maze for ∼30 days^38,39^. All procedures were performed in accordance with the UK Animals Scientific Procedures Act (1986) and approved by the University of Bristol Animal Welfare and Ethical Review Board. We note that this is an archival dataset, originally collected in 2005.

At least 14 days prior to maze training, 16 adjustable tetrodes were targeted to dorsal CA1 of the hippocampus (CA1, 4-6 tetrodes), the prefrontal cortex (PFC, 6 tetrodes), and the parietal cortex (PC, 4-6 tetrodes). Target coordinates were centered on: deep layers of the right prelimbic cortex (+3.2mm, -0.6mm from bregma), CA1 of the right dorsal hippocampus (−4.0mm, +2.2mm from bregma) and deep layers of PC overlying CA1. Custom-built adjustable tetrode (twisted 12.7*µ*m nichrome wire, Kanthal, gold-plated to 250-300kW at 1kHz) microdrives were implanted under isoflurane anaesthesia using aseptic technique and perioperative opiate analgesia (buprenorphine, i.p.). Implants were fixed to the skull using stainless steel screws (M1.4 *×* 2 mm, Newstar Fastenings) and Gentamicin bone cement (DePuy). Tetrode positions were adjusted over the course of 2-3 weeks after surgery.

Signals were amplified by headstages (HS-36, Neuralynx, MT, USA) and relayed via fine-wire tethers to a Digital Lynx system (Neuralynx), which sampled thresholded extracellular action potentials at 32 kHz (filtered at 600-6000Hz) and continuous local field potentials (LFP) at 2kHz (filtered at 0.1-475Hz) using the Cheetah software package (Neuralynx) running on a desktop PC. Differential recordings were made using Local reference tetrodes placed in the superficial prefrontal cortex and in the white matter overlying the hippocampus. Local field potentials (LFPs) originating from each tetrode were bandpass filtered within the frequency range of 0.5 to 475 Hz. Notably, this does not incorporate the full range of location-informative frequencies^29^. The animals’ location on the maze was also tracked using head-mounted LED sampled at 25 Hz. A vizualization of the data collection process is shown in Figure 5.

**Figure 5.**
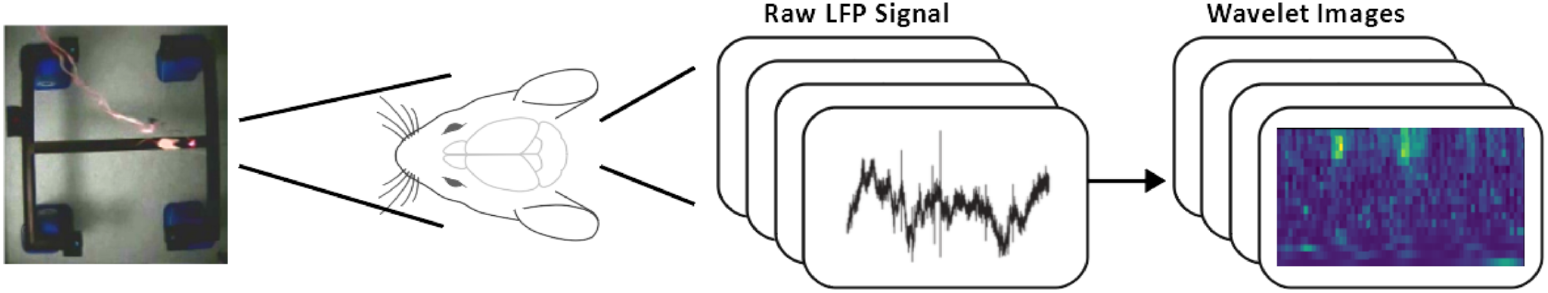
Rats undertook a memory and decision-making task to find sucrose rewards on a maze over ∼35 days. Here we shown a top-down photo of the maze, with rat location tracked via LED on head. Local field potentials were collected from 16 chronically-implanted adjustable tetrodes spanning hippocampal CA1, prefrontal cortex, and parietal cortex. These LFPs were transformed into Morelet wavelet images.

**Figure 6.**
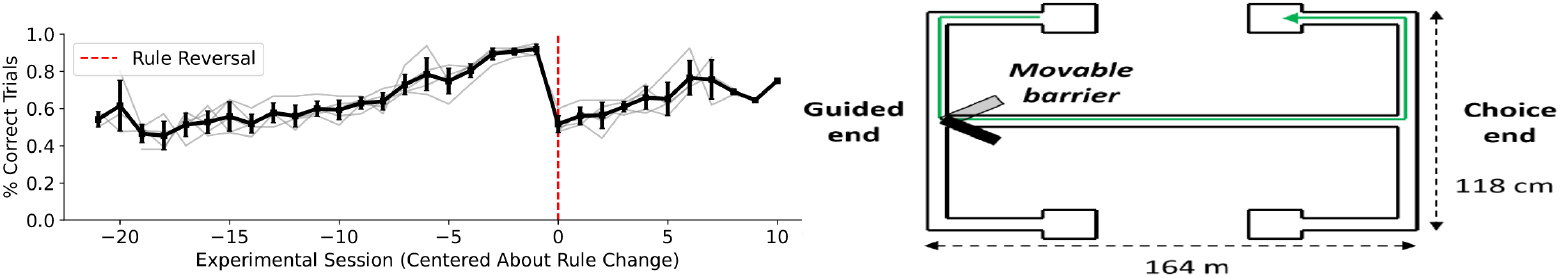
The proportion of experimental trials that were performed correctly for one rat (left). Rats were initially trained under a ‘alternate-turn’ rule - if guided right by barrier, turn left on return opposite end of the maze (right). Once task competency had been achieved, usually ∼25 days, this rule was reversed to a ‘match-turn’ rule.

**Figure 7.**
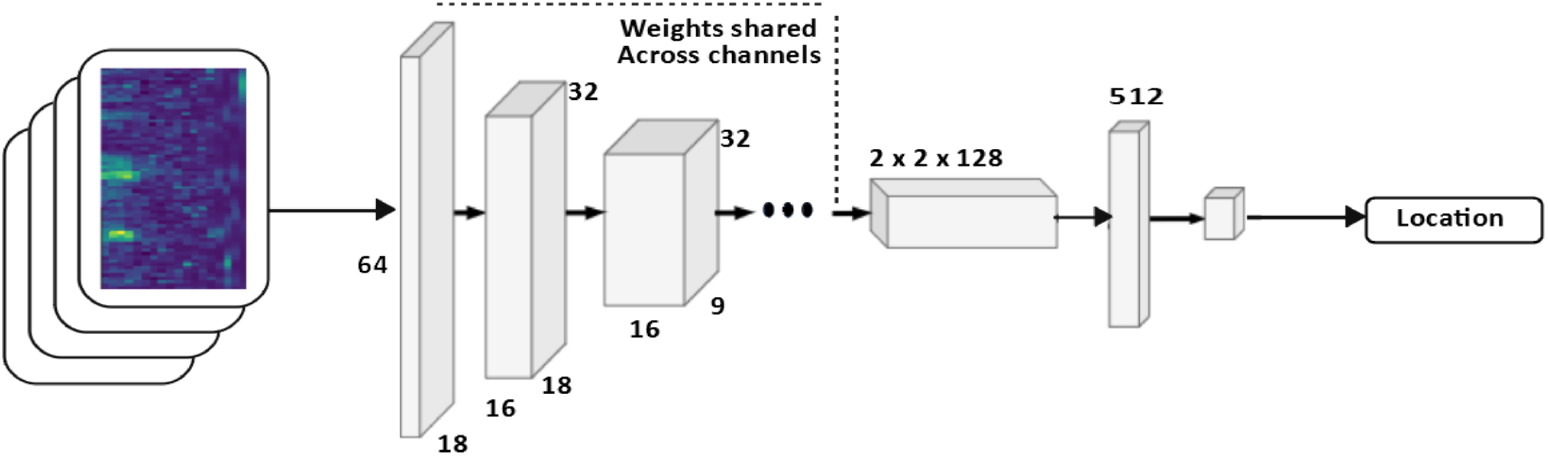
Wavelet images from each experimental session were used to train seperate deep convolutional neural networks which decoded location.

The data collection spanned approximately 30 days, and the rats had minimal prior exposure to the maze, having been introduced to it for less than 1 day before the initial recording. The experimental design required the rats to make choices between left and right maze arms, contingent upon the direction of an initial guided turn. During the initial phase of training, the rats were trained using a “alternating turn” rule, which required that their choices at the choice point oppose their initial guided turn (i.e. if the rats were directed to turn right at the guided turn, they should then turn left at the choice point). When the rats had achieved task competency (three sessions of at least 85% of trials correct), which typically occurred after approximately 20-25 days of training, the rule was reversed to an “match turn” rule, where rats were expected to make choices that were in the same egocentric direction as their initial guided turn (e.g. if they were initially directed right, they were required to turn left at the choice point).

The experimental sessions each day were divided into two distinct phases. The first phase encompassed a series of guided trials, where the rats were guided at both ends of the maze to move in the correct direction using a barrier blocking the incorrect-choice arm. The number of this type of trial varied on a daily basis. Subsequently, the rats underwent a variable number of trials per day, where they were free to make their own decisions at the choice points. Each session, the total number of trials (including guided trials) undertaken by each rat and the proportion of those trials that resulted in successful outcomes was recorded. For 1h before and 1h after maze training, recording continued while rats were in quiet rest and/or sleep states in a sound-attenuating chamber alongside the maze.

LFPs were band-pass filtered at 0.5-475Hz and reflect a spatially weighted aggregate activity of populations of neurons and synapses near the electrode tips^46^. LFP data was then processed into wavelet decomposition images using a Morelet wavelet decomposition^47^ using the PyWavelets package^48^, and wavelet frequencies were downsampled to a rate of 30Hz to discard transient non-local representations. Between experimental sessions, rats were placed in a separate sleeping environment, and hippocampal sharp wave ripples (SWRs) were detected using tetrodes within CA1.

Although the rats were learning task-specific rules during the experiment, the spatial layout of the maze itself remained unchanged throughout data collection. Prior to the experimental sessions, the rats underwent passive exploration to familiarize themselves with the maze’s spatial configuration. During the subsequent sessions, their trajectories consistently covered similar spatial locations, independent of task-specific decision-making. Figure XXX shows histograms of the location distribution of an example animal during sessions 1, 10, and 20; we see little change. This stability in spatial knowledge and location distribution supports the validity of using spatial decoding error to quantify representational drift. By focusing on decoding spatial representations rather than task performance, we isolate changes in neural activity related to oscillatory dynamics and spatial encoding, rather than conflating them with behavioral adaptations or learning of task rules. This ensures that our analysis reflects changes in neural representations of space, not changes in task behavior.

We note that the data used is archival, and has been used in numerous other studies since collection^28,38,39,49,50^.

### Convolutional Neural Network Decoding

In Frey et. al (2021)^29^, a generalizable deep learning framework is presented for decoding sensory and behavioural variables directly from wide-band neural data. The approach requires little user input and generalizes across stimuli, behaviours, brain regions, and recording techniques. This framework has been shown to perform neural localization tasks using Morlet wavelet decompositions of LFP recordings taken from the CA1 pyramidal cell layer in the hippocampus of freely foraging rodents, and is also capable of accurately decoding head direction and speed. We note that this approach to decoding has been shown to be versatile to multiple forms of input including calcium imaging data^51^, thus it is natural to assume that the method presented in this paper would also be versatile to these other modalities.

A modified version of the architecture originally presented in^29^ is used here. Our TensorFlow 2.10.1 implementation uses deep convolutional networks with 13 downsampling time-distributed (i.e. with shared weights across channels) layers followed by two fully connected layers, which output a single (x, y) position estimate for the animal at the time at which the LFP data was collected. A kernel size of 3 was used throughout the model, and the number of filters was kept constant at 64 for the first 10 layers while sharing weights across the channel dimension, then doubling the number of filters for the following 3 layers. For downsampling the input, we used a stride of 2 intermixed between the time and frequency dimension. We used 2D-convolutions which share weights across the channel dimension for the first 10 convolutions and across the time dimension for the last 3 convolutional layers. Sharing weights across channels prevents overfitting to channel-specific features and thus improves generalization by making sure there exists a global representation of important features e.g. of spikes or other prominent oscillations in the local field potential. A visualization of the network is shown in Figure 7, and the full architecture of the network used is presented in Table 1.

**Table 1.** Layer by layer architecture of the convolutional model. Note that for the first 10 layers, weights were shared across the channel dimension, while in layers 11-13, weights are shared across the time dimension. We also note than layers 14-15 exist for each output head (i.e distinct outputs have distinct fully connected layers). Order of dimensions: Channels, Time, Frequency.

| Layer | Layer Type | Input Dimension | Kernel Size | Stride | Filters |
| --- | --- | --- | --- | --- | --- |
| 1 | 2D-Convolution | (16, 64, 18) | (1,3,3) | (1,2,1) | 64 |
| 2 | 2D-Convolution | (16, 32, 18) | (1,3,3) | (1,1,2) | 64 |
| 3 | 2D-Convolution | (16, 32, 9) | (1,3,3) | (1,2,1) | 64 |
| 4 | 2D-Convolution | (16, 16, 9) | (1,3,3) | (1,1,2) | 64 |
| 5 | 2D-Convolution | (16, 16, 5) | (1,3,3) | (1,2,1) | 64 |
| 6 | 2D-Convolution | (16, 8, 5) | (1,3,3) | (1,1,2) | 64 |
| 7 | 2D-Convolution | (16, 8, 3) | (1,3,3) | (1,2,1) | 64 |
| 8 | 2D-Convolution | (16, 4, 3) | (1,3,3) | (1,1,2) | 64 |
| 9 | 2D-Convolution | (16, 4, 2) | (1,3,3) | (1,2,1) | 64 |
| 10 | 2D-Convolution | (16, 2, 2) | (1,3,3) | (1,1,2) | 64 |
| 11 | 2D-Convolution | (16, 2, 1) | (3,3,1) | (2,1,1) | 128 |
| 12 | 2D-Convolution | (8, 2, 1) | (3,3,1) | (2,1,1) | 128 |
| 13 | 2D-Convolution | (4, 2, 1) | (3,3,1) | (2,1,1) | 128 |
| 14 | Fully Connected | (1, 512) | - | - | 1024 |
| 15 | Fully Connected | (1, 1024) | - | - | 1/2 |

Notably, as this system takes minimally-processed input, it is able to perform sensory decoding without spike sorting (a computationally-intensive process for detecting action potentials and assigning them to specific neurons^31^). Necessarily, spike sorting discards information in frequency bands outside of the spike range which potentially introduces biases implicit in the algorithm. Also, as the system is versatile to different forms of input, it is natural to think that the system presented here may also be extendable to non-LFP forms of neural input, for example two-photon calcium imaging data^52^.

Separate networks were trained for each rat using the training portion of the data generated from a single maze task session.

Networks were trained to decode location for 1000 epochs with 250 training steps per epoch. A batch size of 8 was used, as was an AMSGrad optimizer^53^ with a learning rate of 7*e*^−4^.

To quantify performance of our neural decoders, we define error as the Euclidean distance between predicted location and true location, averaged over the test set.

### Random Performance Calculation

To quantify random performance, we trained the networks described in the section above on shuffled input-output pairs, meaning that only the distribution of the output data can be learned by the network. In practice, decoders trained on shuffled data generated outputs close to the mean position of the rat during training on every trial, and generated average decoder error of 84.2cm.

### Drift Quantification

To quantify drift, we first trained the deep convolutional neural networks (CNNs) described above to decode position from wavelet decomposition images of local field potentials (see Figure 7), with distinct networks trained using data from different experimental session.

For each experimental session, separate networks were trained for each rat using data from all brain regions measured (16 tetrodes), as well as three separate networks for each rat trained on data only from CA1 (5 tetrodes), PFC (5 tetrodes) and PC (6 tetrodes). In this way we can examine both local as well as distributed representation dynamics.

Once networks are trained, accuracy data is gathered by calculating the error for networks trained for each day of data and each rat on all other days, generating a table containing error for every train-day/test-day pairing. By calculating decoding accuracy of CNNs trained on data from different days we quantify representational stability of spatial encodings in these regions as a function of train-test-lag. The idea here is that the better a network trained on session *i* performs on session *i* + 1, the more similarities there will be between the spatial representation used by the animal on those days. The end result is, for 30 days of collected data, a 30 *×* 30 accuracy table containing the accuracy scores for all train-test day pairings (see Figure 2).

In order to compare drift rates across animals, and across training and test datasets for the same animal, we normalized the accuracy tables by row (dividing each row’s elements by their sum) to account for model-to-model variability, and then normalized by column to account for test dataset variability.

As our method for drift quantification method is based on the change of decoding accuracy as a function of time, our method can be applied to a variety of sensory or behavioural variables. Drift can be quantified for any representation in any brain region, so long as decoding is possible.

### Drift time constant estimation

The mean normalised accuracy of the tables generated above were well described by exponential decay curves as a function of train-test lag:

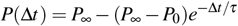

where Δ*t* is the train-test lag, *P*_0_ is the performance of the decoder when training and testing is on data from the same session, *P*_∞_ is the aysymptotic performance of the decoder at very long train-test lags, and *τ* is the drift time constant — the train-test lag at which decoder error will rise by 1*/e* ≈ 63% of the way towards its asymptotic performance.

When pooling the data across training sessions (Figure 2 and Figure 4, we fit the parameters of this function by minimising the squared distance between observed accuracy values and curves using the Levenberg-Marquardt algorithm^54^ in the SciPy 1.10.0 package.

However we found that this algorithm had numerical instabilities when fitting a different time constant per training session (Figure 3). To handle this we used a Bayesian model to perform the session-by-session regressions, which allowed for regularisation of the permitted time constant values. We used the following model:

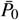∼ Normal(0.5, 0.1)

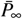∼ Normal(0.6, 0.1)

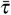∼ TruncatedNormal(5, 5, 1, 30)

*P*_0*i*_ ∼ Normal(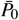, 0.2)

*P*_∞*i*_ ∼ TruncatedNormal(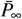, 0.05, 0.5, 0.75)

*τ*_*i*_ ∼ TruncatedNormal(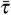, 5, 1., 30)

*σ* ∼ TruncatedNormal(0.1, 0.05, 0.01, 0.3)

*y*(*i*, Δ*t*) ∼ Normal 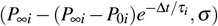 where the bar over the symbol 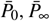 and 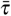 denotes that they are grand means across sessions, *i* indexes the training sessions, *σ* is the error standard deviation and *y*(*i*, Δ*t*) are the actual decoder performance values which here we treat as ‘data’ to be fit. This model performs regularisation via partial pooling of the *P*_0*i*_, *P*_∞*i*_ and *τ*_*i*_ parameters across sessions.

We also used a Bayesian model to perform linear regression on the time constant as a function of training session, pooled across all animals (Figure 3). Bayesian models were fit using the NUTS Hamiltonian monte carlo algorithm in the Turing.jl package^55^ in Julia.

## Statistical Testing

### ANOVA with Tukeys HSD

Statistical testing was undertaken to examine differences between the location decoding error of models trained on data from different brain-regions, between *τ* values generated from these different brain-region models, and between the fit errors of accuracy data ordered by different experimental variables.

ANOVA^56^ was selected as the most appropriate analytical method because it allows for the comparison of the means of three or more independent groups, providing a statistical test of whether or not the means of several groups are all equal. Tukey’s Honestly Significant Difference (HSD) test^57^ was employed as our post hoc analysis method. Tukey’s test is particularly advantageous because it controls for Type I error across all pairwise comparisons, maintaining the overall error rate at the desired significance level. These tests were conducted using the pairwise_tukeyhsd function within the statsmodels 0.13.5 package.

### Ordinary Least Squares Regression

We used ordinary least squares (OLS) regression to obtain best fit linear equations of the location decoder error against experimental session, and of the fitted *τ* values against experimental session. Specifically, OLS minimizes the sum of squared deviations from the actual data points, providing a best-fit linear equation. Using this, we obtained p-values of the slope coefficients to determine statistical significance of the gradient differing from zero. Statistical significance was assessed at the standard p < 0.05 threshold. These tests were conducted using the OLS module within the statsmodels 0.13.5 package.

### Bayesian analysis significance

For the Bayesian analysis of drift time constant as a function of training session (Figure 3), statistical significance was reported as whether the 95% Highest Density Interval for the slope parameter crossed zero). In the text we also report the probability that the slope was below zero, calculated as the fraction of monte carlo samples from the posterior distribution that were negative. Small values for this probability indicate the slope is highly likely to be positive.

## Acknowledgements

We thank Nadine Becker for collecting the data used for this project. We thank the EPSRC (EP/S022937/1), BBSRC (BB/G006687/1, BB/W001845/1), Wellcome Trust (202810/Z/16/Z), MRC (MR/S026630/1) and Leverhulme Trust (RPG- 2019-229) for support. For the purpose of open access, the author has applied a CC BY public copyright licence to any Author Accepted Manuscript version arising from this submission.

